# Plasmid supercoiling decreases during the dark phase in cyanobacteria: a clarification of the interpretation of chloroquine-agarose gels

**DOI:** 10.1101/2021.07.26.453679

**Authors:** Salima Rüdiger, Anne Rediger, Adrian Kölsch, Dennis Dienst, Ilka M. Axmann, Rainer Machné

## Abstract

In cyanobacteria DNA supercoiling varies over the diurnal light/dark cycle and is integrated with the circadian transcription program and (Woelfle et al. [2007], Vijayan et al. [2009], PNAS). Specifically, Woelfle *et al*. have reported that DNA supercoiling of an endogenous plasmid became progressively higher during prolonged dark phases in *Synechococcus elongatus* PCC 7942. This is counterintuitive, since higher levels of negative DNA supercoiling are commonly associated with exponential growth and high metabolic flux. Vijayan *et al*. then have reverted the interpretation of plasmid mobility on agarose gels supplemented with chloroquine diphosphate (CQ), but not further discussed the differences. Here, we set out to clarify this open issue in cyanobacterial DNA supercoiling dynamics. We first re-capitulate Keller’s band counting method (1975, PNAS) using CQ instead of ethidium bromide as the intercalating agent. A 500x–1000x higher CQ concentration is required in the DNA relaxation reaction (topoisomerase I) than in the agarose gel buffer to quench all negative supercoiling of pUC19 extracted from *Escherichia coli*. This is likely due to the dependence of both, the DNA binding affinity of CQ and the induced DNA unwinding angle, on the ionic strength of the buffer. Lower levels of CQ were required to fully relax *in vivo* pUC19 supercoiling than were used by Woelfle *et al*. Next, we analyzed the *in vivo* supercoiling of endogenous plasmids of *Synechocystis* sp. PCC 6803, at two different CQ concentrations. These experiments indicate that negative supercoiling of plasmids does not increase but decreases in the dark phase, and progressively decreases further in prolonged darkness.

## Introduction

*In vivo*, the DNA double helix usually exists in a torsionally strained state, often just denoted as negative DNA supercoiling. The energy stored in underwound DNA can be channeled locally at gene promoters to open and “read” DNA in transcription and replication [Dorman, 2019]. In Bacteria, the level of DNA supercoiling is homeostatically controlled by enzymes but also depends on the transcription and replication, and in turn influences the transcription rates of many genes. In most analyzed bacteria and conditions, increased negative supercoiling is observed during exponential growth and decreased during stationary phase or upon exposure to stress. The DNA supercoiling level of the circular chloroplast genome of the single-celled algae *Chlamydomonas reinhardtii* showed fluctuations with the diurnal light/dark cycle [Salvador et al., 1998]. Mori and Johnson [2001] first suggested that supercoiling could underlie the regulation of hundreds of genes during the light/dark (diurnal) cycle in cyanobacteria, the bacterial ancestors of chloroplasts. Indeed, genome compaction varied over the diurnal cycle in *Synechococcus elongatus* PCC 7942 [Smith and Williams, 2006]. Woelfle et al. [2007] then used an old method, separation of plasmid topoisomers by chloroquine-supplemented agarose gel electrophoresis, to show that the supercoiling level of its endogenous plasmid varies over the light/dark cycle, and variation continued in constant light after entrainment to light/dark cycles. The latter observation is considered a hallmark of the presence of a circadian clock. Specifically, they noted a drop in negative supercoiling of the plasmid already 20 min after the transition to light, as well as continuous increase in DNA supercoiling in prolonged 72 h darkness. However, a comparison with a later paper using the same method [Vijayan et al., 2009] shows discrepancies in the interpretation of the agarose gels, specifically the relative mobilities of more and less supercoiled plasmid samples. We first review the principles of agarose gel electrophoresis of circular DNA in the presence of DNA intercalators. To establish the effect of chloroquine (CQ) on plasmids, we recapitulated Keller’s band counting method but using CQ instead of ethidium bromide (EtBr) as the intercalating agent, and the pUC19 plasmid isolated from *Escherichia coli (E. coli*) culture in exponential growth. Next, we extracted endogenous plasmids from the cyanobacterium *Synechocystis sp*. PCC 6803 (*Synechocystis*) grown under light/dark conditions (12 h/12 h). Using different CQ concentrations, we show that, opposite to the interpration by Woelfle et al. [2007], negative supercoiling increased (lower ΔLk) quickly upon the onset of the light phase and progressivly decreased (higher ΔLk) during a prolonged 48 h dark phase.

## Results and Discussion

### A Short History of Topoisomer Separation

The method to measure plasmid supercoiling by agarose gels is the subject of this contribution and requires some background. Textbook plasmid agarose gels show three distinct bands: the “linear” form which underwent a double strand break, likely during extraction; and the oc (open circular) band, which has only one or more single strand breaks and thus is still circular. The latter travels slowest on the gel. And thirdly, intact plasmids travel fastest as a smeared band, widely known as the ccc band, for the covalently closed circular form of the plasmid. The band is smeared since it actually contains several distinct species, denoted topoisomers, that travel at slightly different speeds in the gel. Topoisomers are identical plasmid molecules (isomers) that differ only by their level of negative DNA supercoiling, quantified as the plasmid’s linking number (Lk) [Crick, 1976]. The terminology is from the mathematical field of topology, and Lk is the number of crossings of the two helix strands around each other. In relaxed (non-supercoiled) DNA the double helix adopts its minimal free energy structure with 10.4 - 10.5 bp per helix turn, *e.g*. a circular DNA with 1000 bp has Lk_0_ = 95 - 96 such crossings. Levels of DNA superocoiling are usually provided as the difference to this relaxed state linking number, ΔLk_0_ = Lk – Lk_0_ and as the supercoiling density 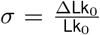. Negatively supercoiled DNA has ΔLk_0_ < 0 and positively supercoiled DNA has ΔLk_0_ > 0. Plasmids isolated from *E. coli* typically have *σ* = −0.08 - −0.06, depending on the growth phase [Liu et al., 2018]. Relative topoisomer abundances follow a Boltzmann distribution around a mean level of *σ* [Pulleyblank et al., 1975, Depew and Wang, 1975, Keller, 1975].

Supercoiled plasmids (`ccc`) form a more compact molecule than relaxed plasmids. This is achieved by a higher level structure where two double helices wind around each other to compensate for the torsional strain on the double helix. This structure, a so-called plectoneme, is the origin of the term supercoiling, since it is a doubly coiled structure: the DNA double helix forms a higher order “double helix”. The more compact plasmids migrate quicker through the polymeric meshwork of an agarose gel. If plasmids are artificially relaxed with a nicking-closing enzyme (NC), such as bacterial topoisomerase I (TopoI), they are still `ccc` but now migrate at a speed similar to the `oc` form. NC enzymes introduce a single strand break (nick). The torsional strain in the supercoiled helix leads to rotation of the single stranded ends. The enzyme then ligates (closes) the ends again and releases a plasmid with less supercoiling. These reactions will result in a Boltzmann distribution of plasmids around ΔLk_0_ = 0, with some positively and some negatively supercoiled [Depew and Wang, 1975]. These latter species are already more compact then the relaxed form and they form distinct bands “below” the oc band.

To separate the *in vivo* levels of supercoiling on agarose gels an intercalating substance can be added to the agarose gel and the gel buffer. Intercalating substances reduce the helix rotation angle between the two bases they bind to, *i.e*., they locally unwind the helix [Lerman, 1961, Pritchard et al., 1966]. This absorbs some of the torsional strain of the underwound helix of negatively supercoiled DNA, and thus reduces the apparent level of supercoiling. When adding the right amount of the intercalator, supercoiling is reduced to an extent which can be separated on the agarose gel. A similar principle can be used to generate a series of plasmid topoisomers, starting from the *in vivo* level of supercoiling and all the way to the fully relaxed form. The intercalator is simply added to the reaction mix, supercoiled plasmids are partially relaxed and the NC enzyme can only remove the remaining level of DNA supercoiling. After washing out the intercalator the formerly quenched supercoiling is re-introduced. Keller [1975] developed an elegant method where different concentrations of the intercalator EtBr are used both in the NC reaction mix and subsequently on agarose gels to separate topoisomers. Keller’s band counting method allows to simply count all bands from the original untreated plasmid to the fully relaxed form across the series of gels, and thereby determine ΔLk_0_ of circular DNA isolated from cell cultures. Importantly, the unwinding by the intercalator is not saturated at full DNA relaxation. Adding more intercalator will result in positively supercoiled plasmids [Shure et al., 1977, Bowater et al., 1992]. These are not affected by typical (ATP-independent) NC enzymes, which only relax negatively supercoiled plasmids [Wang, 1971, Kirkegaard and Wang, 1985]. In contrast, on agarose gels the DNA now changes its migration speed and positively supercoiled plasmids travel again faster than the relaxed form. Shure et al. [1977] suggested that with intercalators with lower affinity the topoisomer separation in agarose gels ‘less *sensitive to variations in experimental conditions*’ and first introduced the use of chloroquine (CQ, all amounts refer to chloroquine diphosphate) instead of EtBr.

### Diurnal Plasmid Supercoiling in Cyanobacteria

Woelfle et al. [2007] used 10 μg mL^−1^ of CQ in their agarose gel analysis. They state but do not show that they have tested the relative mobility changes with different CQ levels, and assume that they are in the range were the plasmids are still negatively supercoiled. That is, plasmids that migrated faster on the gel were interpreted as originally (*in vivo*) more supercoiled (more negative supercoiling, lower Lk). This is reflected in their gel band annotation and the equation they used to calculate an average relative mobility (*RM*) of each sample:

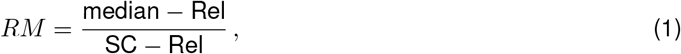

where “Rel” denotes the migration distance of the upper (slowest migrating) band, actually the still present oc form. A lower, fastest migrating band that also appeared in all samples is denoted as “SC”. However, it is unclear what this band actually is, because the supercoiling is quenched by CQ and the actual supercoiled plasmids travelled as multiple topoisomers. A “median” of topoisomer migration speed was used to calculate the *RM* of each sample. Subsequently, Vijayan et al. [2009] used the same species, the same endogenous plasmid and the same principle to analyze plasmid supercoiling after gyrase inhibition and correlate it to global changes of the transcriptome. They only tested constant light conditions. However, they have subtly changed the protocol and used 15 μg mL^−1^ of CQ in the agarose gels. Without any discussion of this issue they have further reverted the interpretation of migration speed to:

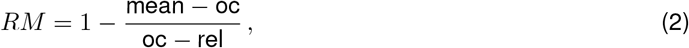

where the annotated gels indicate that they inpret the upper, slowest migrating band as the `oc` form. They also observe a fast migrating band in all samples which for unexplained reasons they denote as a “rel” for relaxed. They calculated a “mean” migration distance of the supercoiled topoisomers. The equation likely contains an error, since *oc* – *rel* would be a negative value. Independent of these unclarities the subtract the *RM* of Woelfle *et al*. from 1. This implies that they interpret faster migrating topoisomers as originally less negatively supercoiled. At high intercalator concentration these should be more positively supercoiled and travel faster. Choosing more neutral band identifiers makes these differences clearer:

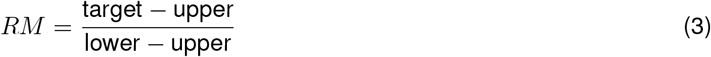

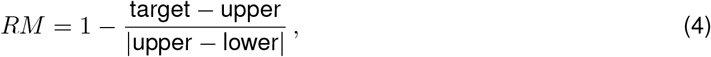

where “upper” and “lower” refers to their position on the original gel images, and “target” is the mean or median of the topoisomer distribution. Despite the opposite meaning of *RM*, both publications interpret a higher *RM* as a higher level of negative DNA supercoiling.

This different interpretation of the gels in Woelfle et al. [2007] would imply that negative supercoiling of plasmids rapidly increases upon onset of the light phase and would progressively decrease in prolonged darkness. Vijayan et al. [2009] measured supercoiling only in constant light conditions, and did not mention or discuss these discrepancies. Thus, the DNA supercoiling dynamics within the natural light/dark cycle of cyanobacteria remains unclear.

### Keller’s Band Counting Method with Chloroquine and pUC19

[Keller, 1975] analyzed the linking number of the circular genome of the simian virus 40 (SV40, 5.2 kb), propagated in and isolated from African green monkey cells (CV-1). For this, the DNA was subjected to *in vitro* relaxation with an NC enzyme purified from human tissue culture cells (KB-3). Full relaxation of the SV40 DNA in the gel buffer was achieved at *C*_gel_ = 0.06 μg mL^−1^, or with 394.3 g mol^−1^, *C*_gel_ = 0.15 μM. In the topo I reaction *C*_relax_ ≈ 4.5 μM (lane 9 in Figures 1 and 3 of [Keller, 1975]) were required to fully relax the DNA. The *in vivo* supercoiling level of the SV40 DNA was ΔLk_0_ ≈ 24 and *σ* ≈ 0.05.

**Figure 1.**
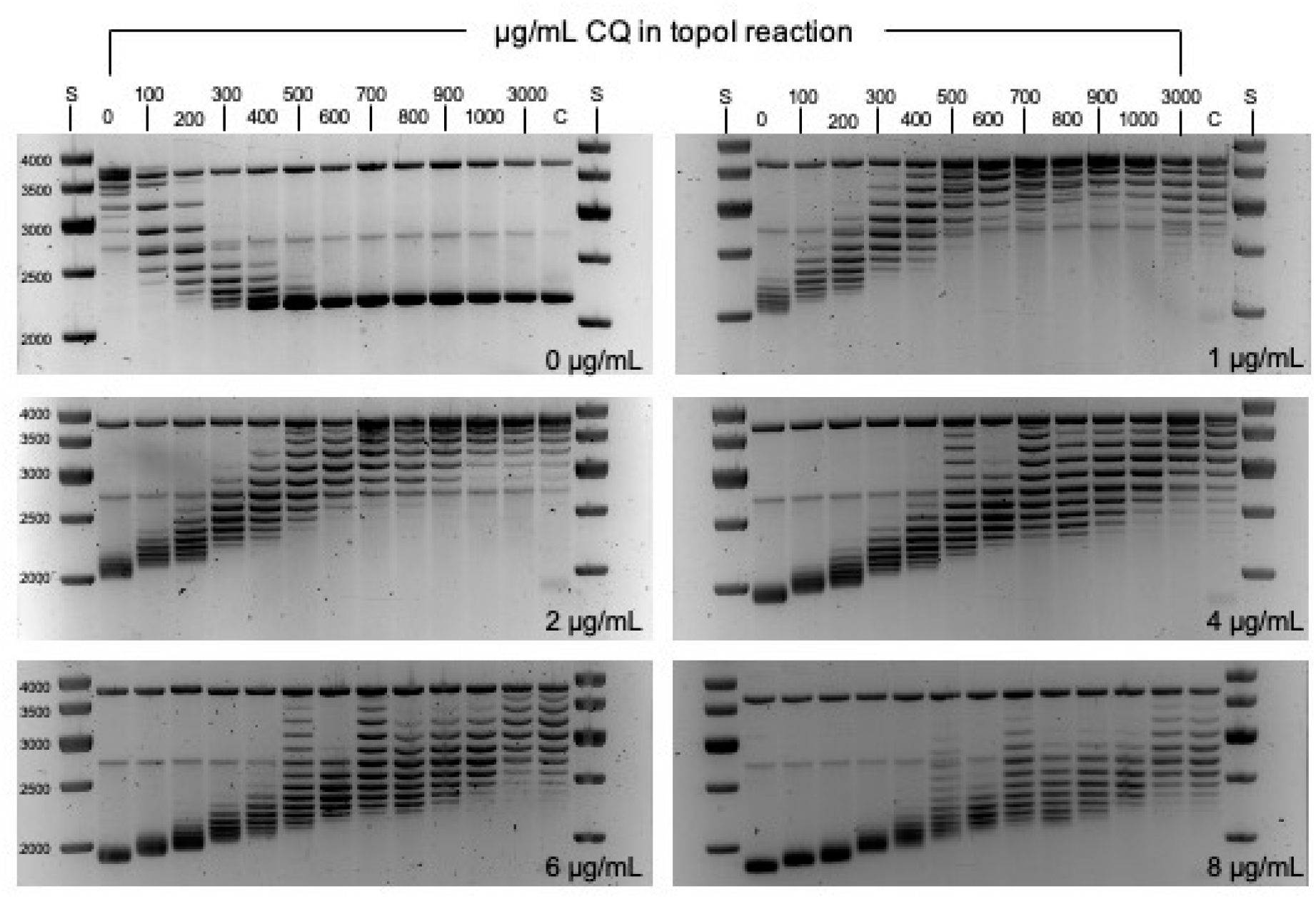
pUC19 on 1.2 % agarose gel supplemented CQ. Samples of the pUC19 plasmid, isolated from *E. coli*, were partially relaxed with *C*_relax_ = 0 - 3000 μg mL^−1^ CQ and topoI as indicated (lanes). The sample “C” is untreated pUC19 as control. Size marker “S” is the GeneRuler 1 kb DNA Ladder. The samples analyzed on 6 agarose gels, each supplemented with a CQ concentration (*C*_gel_ = 0 - 8 μg mL^−1^) as indicated (lower right). The top band in each gel is the open circular form. The band between 2.5 kb and 3 kb is the linear form of the plasmid.

Here, we propagated the pUC19 vector plasmid (2686 bp) in in *E. coli* DH5*α* and harvested plasmids in exponential phase on LB medium. A plasmid relaxation series was performed with the bacterial NC enzyme TopoI obtained from New England Biolabs. To find the appropriate CQ concentrations to reproduce Keller’s band counting method, we first analyzed the EtBr concentrations used in the original publication, considered the ca. 1000x lower binding affinity of CQ, and adjusted concentrations from there. Figure 1 shows that the lowest CQ concentration in the gel (*C*_gel_ = 1 μg mL^−1^) was sufficient to detect a clear change in the migration of all pUC19 topoisomers. Fully relaxed plasmids (*C*_relax_ = 0 - 200 μg mL^−1^) had already shifted to positive supercoiling and migrated faster than without CQ. Topoisomers treated with *C*_relax_ = 800 - 1000 μg mL^−1^ were all close to full relaxation at *C*_gel_ = 1 μg mL^−1^. The untreated *in vivo* sample (“C”) appeared fully relaxed at *C*_gel_ = 2 μg mL^−1^, and shifted to positive supercoiling at *C*_gel_ > 4μg mL^−1^. The topoisomers relaxed at the highest concentration (*C*_relax_ = 3000 μg mL^−1^) appeared with band patterns almost identical to the untreated control in all gels. With a molecular weight of 515.86 g mol^−1^ (chloroquine diphosphate), fully relaxing concentrations of CQ were thus *C*_gel_ ~ 3.9 μM and *C*_relax_ ~ 5.8 mM. Counting topoisomers across gels shows that the untreated control has a ΔLk_0_ = −15 - −16. With 2686 bp length, Lk_0_ ≈ 257 and thus *σ* ≈ 0.06, as expected for a plasmmid isolated from *E. coli*.

In summary, the intercalator concentrations required to fully quench the original level of supercoiling of SV40 and pUC19 were ≈30x and ≈1000x higher in the reaction buffers than in the gels, respectively. The CQ concentrations were 100 - 1000x higher than the EtBr concentrations in Keller [1975]. EtBr binds to DNA with relatively high affinity, *e.g*. *K_a_* = 1.5 × 10^5^ M^−1^ in 0.2 M Na^+^ [Gaugain et al., 1978], and each intercalated molecule unwinds the helix by 26 - 28° [Keller, 1975]. CQ has a lower affinity to DNA, depending on ionic strength of the buffer, *e.g*. *K_a_* = 3.7 × 10^4^ M^−1^ in a 50 mM phosphate buffer and *K_a_* = 3.8 × 10^2^ M^−1^ with 0.1 M NaCl [Kwakye-Berko and Meshnick, 1989]. The DNA unwinding angle induced by CQ is lower and, unlike EtBr, also depends on the ionic strength of the buffer, with a maximum of 17° at 0.05 M salt concentration, and CQ likely has additional non-intercalative binding modes [Jones et al., 1980]. Additionally, intercalator effects as well as supercoiling-dependent conformations of circular DNA could be sequence dependent [Vetcher et al.,2010]. Likely, all these factors contribute to the differences in the concentrations required for full relaxation of SV40 by EtBr, of pUC19 by CQ, and between the gel and reaction buffers in both implementations of Keller’s band counting method.

### Gradual Plasmid Relaxation in Dark Phase, Quick Supercoiling in Light

To clarify the unresolved issue whether DNA supercoiling is higher during the light or dark phases in cyanobacteria, we measured supercoiling levels of endogenous plasmids of *Synechocystis sp*. PCC6803 (“Moscow strain” PCC-M) during diurnal light/dark (12 h/12 h) conditions and a prolonged dark phase (12 h light, followed by h24h darkness). A large sample volume (25 mL) was required to obtain enough plasmids, and direct mixing of the sample with an equal volume of pre-cooled (−20°C) pure undenatured ethanol was key to successful plasmid extraction. However, the yield was still very low (0.5 - 1 μg, but incl. genomic DNA), limiting the amount of gels that can be run from one sample. Multiple sets of putative topoisomer bands were detected at all sampled time points by agarose gel electrophoresis supplemented with chloroquine at either 1μg mL^−1^ or 20 μg mL^−1^ (Fig. 2A, B). *Synechocystis* contains three short plasmids, two at 2.4kb [Yang and McFadden, 1993, 1994] and one at 5.2 kb [Xu and McFadden, 1997]. Restriction analysis and Southern blots confirmed that the most pronounced topoisomer bands were obtained for plasmid pCA2.4_M. The migration distances on the gels were different between samples and the relative differences reversed between the two CQ concentrations. This change of relative migration speed indicates that plasmids are still negatively supercoiled at the low, but positively supercoiled at the high concentration. Plasmids which were originally more relaxed migrate slower and faster at the low and high concentrations, respectively. Next, we quantified the agarose gels by generating electropherograms of each lane in ImageJ. Baseline correction, peak detection and peak area quantification were performed in R (Fig. 2C). For each sample an average linking number deficit 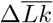 was calculated and plotted as time series over the light/dark cycles (Fig. 2D). Plasmids were more negatively supercoiled (lower 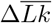) during light phases, and reached a maximal level only 30 min after transition to the light phase. This level was maintained or only slightly increased throughout the 12 h light phase. During the dark phase the transition was slower, and plasmids became progressively more relaxed during the 12 h, and relaxation continued at equal pace during an additional 12 h dark phase. The maximal linking number differences between light and dark phase were only 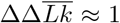, and 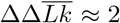 after the prolonged dark phase.

**Figure 2.**
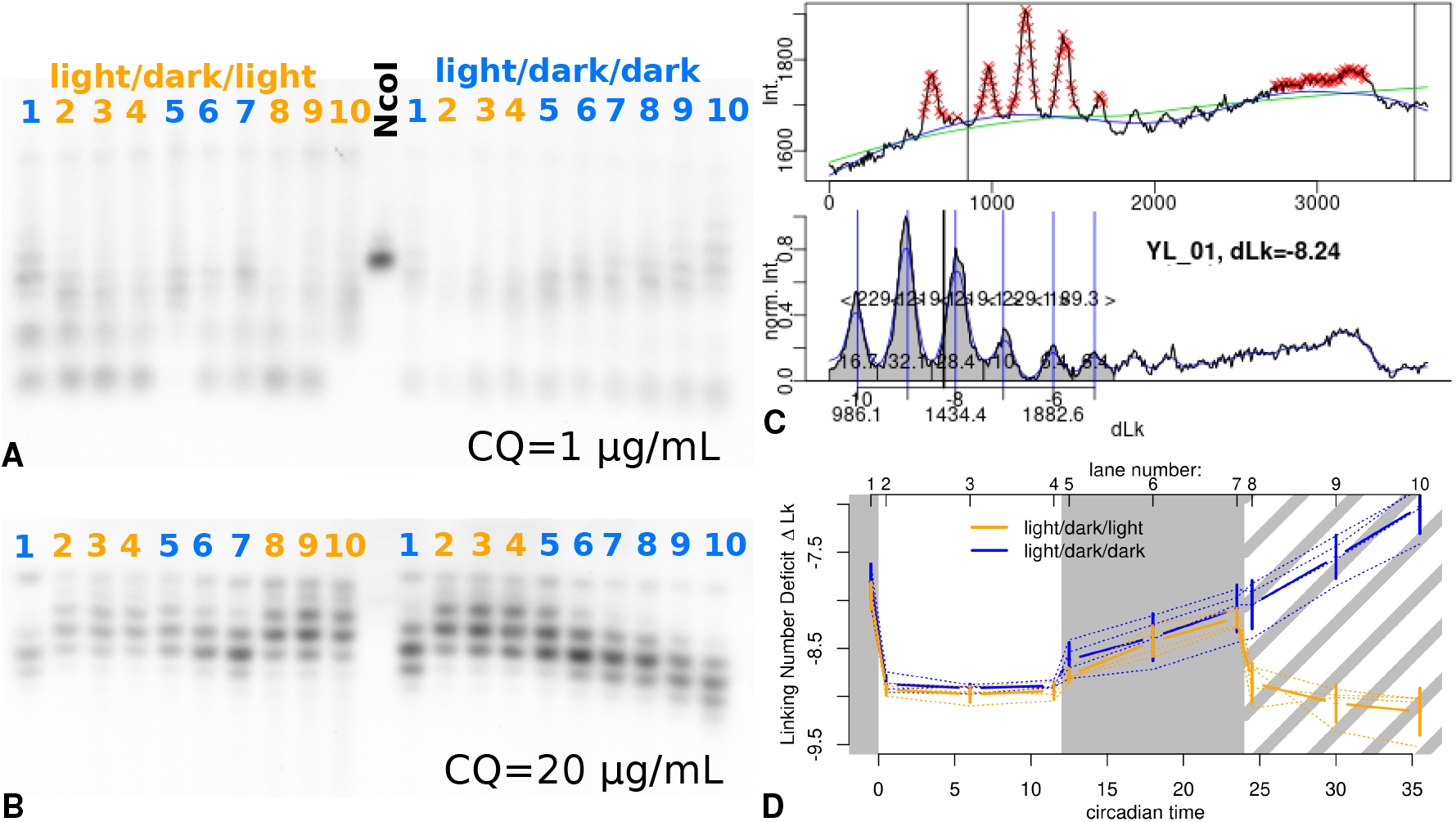
Diurnal Plasmid Supercoiling in *Synechocystis*. **A & B:** Southern blots with a probe for the pCA2.4_M plasmid of 1.2%agarose gels supplemented with 1 μg mL^−1^ (A) or 20μg mL^−1^ (B) CQ. The samples were extracted from two diurnal growth experiments and applied in the same order on both gels, see top axis in (D) for sampling times of the lanes. The central lane in (A) shows a pooled plasmid sample after treatment with the NcoI restriction enzyme which cuts only the pCA2.4_M plasmid (one cut site) but not the similarly sized pCB2.4_M. **C:** Example electropherogram of a single lane of topoisomers, extraced with ImageJ from a gel image and analyzed in R using peak detection routines to define a baseline in two steps (top panel, blue and green lines) and to calculate a mean linking number from topoisomer peak areas (bottom panel, 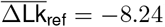). **D:** Diurnal time series of the average linking number deficits of the pCA2.4_M plasmids, calculated from three replicates of agarose gels with 20 μg mL^−1^ (B) and the single replicate of the gel with 1 μg mL^−1^ (A). The top x-axis maps samples to the lanes in (A) and (B). The dotted thin lines are values from the 4 different gels and the thick lines are their means and standard deviations. Gray background indicates the dark phases. One culture (blue line, light/dark/dark) did not receive the final 12 h of light and remained in the dark. Note, that lower values indicate more negative supercoiling (lower 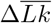). The absolute values of ΔLk are an estimate of ΔLk_0_, based on the distance to the relaxed forms after topoI relaxation (not shown), and likely underestimates the real |ΔLk_0_|.

In summary, negative supercoiling of pCA is higher during the day, with a quick increase directly after the onset of light, and lower during the night; extended night relaxes plasmid even more. These results are the exact opposite of the results reported by Woelfle et al. [2007] for the endogenous plasmid of *Synechococcus elongatus* PCC 7942. We believe that Woelfle et al. [2007] had misinterpreted the relative topoisomer mobilities, and the results are actually consistent between both species. This conclusion is further supported by the reversal of the interpretation of relative topoisomer mobilities by Vijayan et al. [2009].

## Conclusion

Plasmid topoisomer analysis by intercalator-supplemented relaxation series and agarose gel electrophoresis is an old and elegant method. However, the gels are difficult to interpret, and the effect of intercalator concentrations must be checked for each plasmid and conditions (gel and reaction buffers). Ideally, Keller’s band counting method should be applied to each newly analysed plasmid. The low yields when extracting endogenous plasmids from cyanobacteria, the time-intensive calibration routine, and non-standard electropherogram analysis make it somewhat challenging to properly analyse *in vivo* supercoiling in a given species.

Can these methods be updated to the requirements of current day laboratory practice? Mitchenall et al. [2018] reported separation of topoisomers of pBR322, pUC19, and a 339 bp DNA minicircle on a commercial capillary gel electrophoresis platform (Qiagen QIAxcel Advance System), where EtBr is a component of the gel and used by the detection system for quantification. The system is quite sensitive and samples of 100 ng DNA sufficed. Moreover, the range of separated topoisomers was broader than on agarose gels. This would in principle allow to use fewer runs to achieve full separation of *in vivo* levels of supercoiling. However, the intercalator type and concentration can not be manipulated in this gel cartridge-based system. We recently succeeded to separate pUC19 topoisomers by adding comparatively large concentrations of CQ (*C*_gel_ = 1 - 3 mg L^−1^) to the commercial (and secret) gel delivered for the AATI Fragment Analyzer (now owned by Agilent). But the results were not consistent between runs. Likely, the undisclosed fluorescent nucleic acid marker has effects on DNA supercoiling. More work and alternatives to the existing commercial systems are required to establish topoisomer separation by capillary gel electrophoresis as a standard procedure. This could potentially greatly reduce the amount of sample required per run.

The analysis of electropherograms is also a limiting factor for many laboratories. No standards or established tools exist on how to calculate average linking number differences between samples. Commerical gel image analysis software is not geared towards this application. To this end Ziraldo et al. [2019] provide a plug-in for the ImageJ image analysis software that could streamline the analysis of agarose gels. Here, we also used ImageJ to extract electropherograms from gel images, and peak detection routines from a mass-spectrometry R package, `msProcess` (see Methods) which is not maintained anymore, to calculate average linking numbers in R. A novel R package, `bioanalyzeR` (https://github.com/jwfoley/bioanalyzeR), allows to parse electropherograms from data files exported by Agilent capillary electrophoresis platforms (Bioanalyzer, TapeStation). And finally, Vetcher et al. [2010] recently suggested a model that can partially explain topoisomer mobility in agarose gels, based on the writhe component of ΔLk. Implementing such models in analysis tools could further support automatization. Standardized and automated analysis pipelines for both, gel images and capillary electrophoresis platforms, could open this old and elegant method for modern molecular biology laboratories.

## Data Availability

The topoI relaxation series and agarose gel protocols are available online at `protocols.io`, see Methods section. Images, electropherograms and custom R analysis scripts are available on request.

## Funding

RM was funded by the *Deutsche Forschungsgemeinschaft* (DFG), grants AX 84/4-1 & STA 850/30-1 (COILseq) and EXC-2048/1–project ID 390686111 (CEPLAS).

## Materials and Methods

### pUC19 Plasmid Extraction

The vector DNA plasmid pUC19 (Roth: X911.1) was transferred into *Escherichia coli* DH5*α* via heat shock transformation and clones selected on LB plates with the pUC19 selection marker (100 μg mL^−1^ ampicillin). A single colony was inoculated into LB medium (+ampicillin) and grown overnight at 37°C and 250 rpm. The preculture was diluted 1:200 in fresh LB (+ampicillin) and harvested in exponential phase after 3 h. The culture was centrifuged and pUC19 was isolated with ZymoPURE’s Plasmid Maxiprep Kit. The isolated plasmid was then purified (linear and open circular forms digested) via T5 exonuclease (NEB: M0363) reaction and the NucleoSpin Gel and PCR Clean-up kit (Machery-Nagel). The final plasmid DNA concentration was determined with the Nanodrop (Thermo Scientific NanoDrop 2000c).

### pUC19 Relaxation Series

The pUC19 plasmid extract was split and 1 μg DNA was mixed with 0 - 3000 μg mL^−^ (final concentration) of chloroquine diphosphate salt (CQ, Sigma: C6628-50G, CAS: 50-63-5), 2.5 μL 10x CutSmart reaction buffer (NEB: B7204, 500 mM potassium acetate, 200 mM Tris-acetate, 100 mM magnesium acetate, 1000 μg mL^−1^ BSA, pH 7.9), filled up to 24 μL with nuclease-free water, and incubated for 15min at 37°C to allow for equilibration of the CQ intercalation. Then 1 μL (5U) topoI (NEB: M0301) was added and the reaction mix incubated for another 15 min at 37 °C. Then, the reaction was stopped by incubation at 65 °C for 20 min and samples were purified with the NucleoSpin Gel and PCR Clean-up kit (Machery-Nagel) to remove the enzyme and the intercalator. This protocol is published at https://dx.doi.org/10.17504/protocols.io.rbcd2iw.

### *Synechocystis* Strain and Culturing Conditions

Diurnal plasmid supercoiling time-series were established at conditions and time-points identical to those used for the transcriptome study in ref. [Lehmann et al., 2013, Beck et al., 2014]. The glucose-tolerant and motile wild-type strain PCC-M of *Synechocystis* sp. PCC 6803 (obtained from S. Shestakov, Moscow State University, Russia), was grown photoautotrophically in BG11-medium at 30 °C under continuous illumination with white light at 80 μmol m^−2^s^−1^ (Versatile environmental test chamber; Sanyo) and with a continuous stream of air in two glass tube reactors with 800 mL culture volume. The optical density at 750 nm of the culture was monitored (Specord200 Plus; Analytik Jena). Cultures where then entrained to 12 h/12 h light/dark cycles for three consecutive days and diluted to OD_750_ ≈ 0.5 one day before sampling. Samples for plasmid analysis and OD_750_ were taken at the indicated timepoints for 1.5 days. One culture was kept in dark for the last 12 h of sampling.

### Plasmid Extraction from *Synechocystis* Cultures

25 mL of cell culture were mixed with 25 mL of pre-cooled undenatured 95% ethanol (−80°C and on dry ice during sampling), in 50mL centrifuge tubes and stored at −80 °C until processing. After thawing on ice, the supernatant was discarded after centrifugation for 10 min at 4 °C and 4000 g. The QIAprep Spin miniprep kit was used according to manufacturer’s instruction, except for additional enzymatic steps during lysis. The cell pellet was resuspended in 250 μL Qiagen P1 solution and transferred to 1.5 mL reaction tubes. Then 50 μL lysozyme solution (50 mg mL^−1^) was added, mixed, and incubated for 1h at 37 °C. After the addition of 55 μL of 20 %SDS and 3 μL of proteinase K (20 mg mL^−1^), the reaction mixture was incubated at 37 °C for 16 h. Starting with the alkaline lysis with the Qiagen P2 solution, all further steps (QIAprep Spin Miniprep Kit) were carried out with amounts that were adjusted to the initial volume. Next, the concentrations and quality (260/280, 230/280 ratio) was determined using the Nanodrop (Thermo Scientific NanoDrop 2000c). 30 μL of the extracts (1 - 2 μg of DNA) were digested with PlasmidSafe enzyme mix (epicentre, cat. no. E3101K) to remove linear DNA according to the manufacturer’s protocol and incubated at 37 °C for 30 min and this reaction mix was directly loaded into agarose gel wells.

### *Synechocystis* Plasmid Restriction Analyses

A pooled sample of plasmid extracts from *Synechocystis* was subjected to rectriction by NcoI (Thermo Scientific: FD0573), which has a single cut site in pCA2.4_M but not in the similarly sized pCB2.4_M. 1.5 μL of FastDigest buffer and 1 μL of restriction enzyme were added to 12.5 μL of plasmid samples. Reactions were incubated for 30 min at 37 °C and stopped by addition of 3 μL of 6x DNA loading dye with 1 %SDS and incubation at 80 °C for 15 min. The resulting 18 μL were loaded directly onto the agaorse gels

### Chloroquine Agarose Gel Electrophoresis

Agarose gels with different concentrations of chloroquine diphosphate salt (CQ, Sigma: C6628-50G, CAS: 50-63-5) were used to determine the relative migration speed of supercoiled topoisomers. 1.2% agarose gels in 0.5x TBE (Roth: 3061.2; 0.5x: 5.4 g/L Tris base, 2.75 g/L boric acid, 4 mL/L of 0.5 M EDTA (pH 8.0), pH 8.3) were prepared by heating to boiling. After cooling (hand-warm) CQ was added to the indicated final concentration from a stock solution (10 mg mL^−1^) and the mixture poured into the gel chamber. The running buffer was 0.5x TBE buffer with the same CQ concentration as the gel. For each sample, 250 ng (*E. coli*/pUC19 after treatment with T5 exonuclease) or 1 - 2 μg (*Synechocystis*, PlasmidSafe reaction mix was directly loaded) of plasmid DNA were mixed with loading dye and filled up to 30 μL with water. Gels were run for 16 - 24h at 40 V (1.8 V cm^−1^) in a Peqlab gel chamber, covered in aluminum foil to protect from light. Gels were then washed two times for 30 min in 250 mL 0.5x TBE buffer to remove the CQ, and stained with 25 μL Sybr Gold in 225 mL 0.5x TBE buffer for 3 - 24 h and imaged on a BioRad Imaging System (ChemiDoc MP). Washing and staining were also performed light-protected. The protocol is published at https://dx.doi.org/10.17504/protocols.io.nbtdann.

### Southern Blots of Agarose Gels

Probes for Southern blots of the pCA2.4_M plasmid were generated by colony PCR, using forward primer: ACAGGGGTAAATGAGTGCCG, reverse: GCAAGCAGTCCTCCACAAGA, and the PCR programm: 3 min, 95 °C; 35x(95 °C 30s, 50 °C 30s, 72 °C 1 min); 72 °C 5 min. The products were isolated by cutting the band from an agarose gel and clean-up with the NucleoSpin Extract II kit (Macherey-Nagel). Labelled DNA was generated using the same PCR program with the DIG Easy Hyb (Roche, 1603558) mix, according to manufacturer’s instruction. Southern blots were generated using the CDP-Star kit with slight modifications as follows. Gels were blotted onto a nitrocellulose membrane using a vacuum blotter (90 min with 5 - 7 mmHg, in 10x SSC buffer). The membrane was pre-hybridized for 1 h at 50°C. The probes were denatured (95 °C, 15 min) and cooled on ice, then transfered to the blot and hybridized over night at 50 °C. The membran was washed 2x 5 min mit 2x SSC, 0.1 % SDS at 50 °C, then with 2x 15 min mit 0.1x SSC und 0.1 % SDS at 65 °C, and cross-linked with UV light for 10 min. All further steps were performed according to the CDP-Star manual and blots were imaged on a BioRad Imaging System (ChemiDoc MP).

### Analysis of Gel Electropherograms & Calculation of Δ*Lk*

Electropherograms were extracted in ImageJ for each lane and analyzed in R, using LOESS smoothing and peak detection functions from the msProcess R package (version 1.0.7) (https://cran.r-project.org/web/packages/msProcess/). A baseline was determined in two steps using the `msSmoothLoess` function (Fig. 2C, top panel). The first step used the full signal and served to determine the coarse positions of peaks. The final baseline was then calculated from the signal after removal of peak values. This baseline was subtracted from the total signal to detect peaks (bands) with the `msPeakSimple` function from `msProcess` and calculate peak areas. For topoisomer analysis peaks were detected with the `msPeakSimple` function and the assigned consecutive integral linking number values Δ*Lk_i_* with an estimated offset of a reference peak from the relaxed form with *Lk* = 0. The areas under the peaks *A*_Δ*Lk*_ were calculated and the average linking number deficit of a sample was then determined as the center of mass

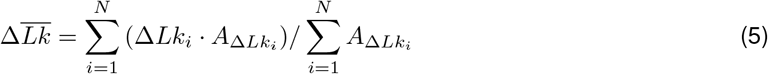

of all *N* identified topoisomer bands.

Note, that the true Δ*Lk* of bands from the relaxed plasmid was not determined, eg. by Keller’s band counting method [Keller, 1975]. The calculated Δ*Lk* values refer to an assignment of a reference band that was present in all lanes to Δ*Lk* = −8, and this can be considered a minimal estimate of the true |Δ*Lk*|, based on the distance of topoisomer and relaxed DNA bands in a topoI relaxation experiment (not shown).

## Notes

### Competing Interest Statement

The authors have declared no competing interest.

